# Representational similarity analysis of EEG reveals multiple spatiotemporal dynamics of selective attention

**DOI:** 10.64898/2026.06.18.733142

**Authors:** Jinhee Kim, Winko W. An, Abigail Noyce, Barbara Shinn-Cunningham

**Affiliations:** Department of Psychology, Carnegie Mellon University, Pittsburgh, PA, USA; Developmental Medicine, Boston Children’s Hospital, Boston, MA, USA; Harvard Medical School, Boston, MA, USA; Neuroscience Institute, Carnegie Mellon University, Pittsburgh, PA, USA

## Abstract

Selective attention allows listeners to follow a target speaker in complex scenes, but less is known about the neural mechanisms that control this capacity. We collected electroencephalography (EEG) during an auditory attention task encompassing spatial attention, talker-based attention, and passive listening. Time-resolved representational similarity analysis (RSA) of broadband scalp voltage and alpha-band oscillations revealed distinct representational trajectories in the two measurements: scalp voltage produced brief peaks in task discriminability shortly after the target cue and around 300 ms after target onset, whereas alpha-band power emerged more slowly and remained elevated throughout the trial. These patterns distinguish attentive from passive conditions and spatial from talker attention. Furthermore, broadband activity and alpha power encode complementary information, while other frequency bands contribute little independent explanatory power. These results demonstrate that RSA allows a unified interpretation of evoked and induced EEG measures, revealing how distinct neural computations jointly encode the spatiotemporal dynamics of selective attention.

## 1. Introduction

Whether in a busy restaurant trying to order food or attending a Zoom meeting in a cafe, we face the daily challenge of selecting and focusing on one auditory target among many competing sounds. This challenge, known as the “cocktail party problem” (Cherry, 1953), highlights the importance of auditory selective attention—the ability to focus on a specific sound source while ignoring distractions (Alain & Bernstein, 2008). Understanding how auditory selective attention is implemented and controlled is critical to explaining how we perceive speech in noisy environments. Further, because even modest hearing impairment interferes with auditory selective attention (Dai et al., 2018), identifying the mechanisms that support selective listening in noisy environments has substantial clinical relevance.

Listeners rely on a range of cues to identify and selectively attend to task-relevant speech amidst distractors. One prominent and well-studied cue is talker location. The representation of the location of a sound source first emerges in the brainstem and propagates to auditory cortex; when behaviorally relevant, spatial position can even be decoded in frontal cortex (Blauert, 1997; Lewald & Getzmann, 2011; van der Heijden et al., 2019; Middlebrooks, 2021). Another key cue is the voice identity of the speaker, shaped by fundamental acoustic features such as pitch and timbre (Darwin et al., 2003; Oh et al., 2021; Shinn-Cunningham, 2008). Differences in fundamental frequency and vocal tract length allow listeners to distinguish individual talkers even in challenging listening conditions (Darwin et al., 2003). In addition, the structure of the speech signal itself contributes to attentional selection. Temporal features in the speech envelope help listeners track a continuous speech stream (Henry & Obleser, 2012; Obleser & Kayser 2019). While listeners are able to use these different types of cues to focus on task-relevant speech, the underlying neural mechanisms through which attention employs each cue remain only partially understood.

Electroencephalography (EEG) provides a powerful tool for studying neural activity during auditory attention, primarily due to its high temporal resolution (Näätänen & Picton, 1987; Luck, 2014). Attention modulates neural activity through multiple mechanisms, and EEG recordings reflect these modulations (Tavano et al., 2023). For example, auditory event-related potentials (ERPs) are modulated by attention when listeners track one of two competing streams (Choi et al., 2013; Hillyard et al., 1973; Nguyen et al., 2024). Frequency-based features also reflect attentional modulation. Alpha-band power (8-12 Hz) in parietal regions is a hallmark of spatial attention (Kerlin et al., 2010; Wöstmann et al., 2021). Parietal alpha lateralization correlates with spatial cue direction—alpha power increases ipsilateral to the attended location and decreases contralaterally (Deng et al., 2020). Parietal alpha power also has been implicated in non-spatial auditory attention (Wöstmann et al., 2017; Viswanathan et al., 2023). Neural activity in lower frequency bands like delta (2-4 Hz) and theta (4-8 Hz) may also be important in selective processing of attended speech, as they closely track speech features such as temporal envelope (Etard & Reichenbach, 2019; Ding et al., 2016).

Despite these decades of studies, it remains unclear how various EEG features index the neural mechanisms supporting different aspects of auditory attention. There are a number of challenges. First, the high dimensionality of EEG data makes it challenging to compare condition-related differences in a single view. Different types of auditory attention lead to different patterns of coordinated activity across multiple, spatially distributed brain regions. Decoding these differences requires fine-grained temporal resolution and access to the spatial distribution of neural activity across various EEG features, such as time-domain signals and frequency band power. Second, comparing these EEG features is challenging because they differ in both spatial and temporal characteristics. For instance, ERPs reflect short-latency, stimulus-evoked activity in frontocentral regions, while alpha-band oscillations reflect sustained spatial attention in posterior areas. Third, the magnitude of these features differ, complicating efforts to combine information across them.

Representational similarity analysis (RSA; Kriegeskorte et al., 2008) offers a unified framework for describing neural activity across conditions and measurement techniques. The approach first computes pairwise similarity between conditions and then infers their representational structure from the resulting pattern of all comparisons. From its initial applications in examining visual representations in the brain using fMRI (Kriegeskorte et al., 2008; Cichy et al., 2014), RSA has since been applied to fMRI data to explore diverse topics such as semantic knowledge and episodic memory (Clarke & Tyler, 2014; Xue, 2018). More recently, RSA has been used to exploit fine-scale temporal information of EEG signals to gain insights into object recognition, audiovisual processing, and reading comprehension (Kaneshiro et al., 2015; Cecere et al., 2017; Wang et al., 2020). For example, Cecere et al. (2017) computed time-resolved neural representational patterns to assess the similarity between auditory-leading and visual-leading multimodal stimuli, showing that they elicit distinct neural representational structures.

A key advantage of RSA emerges when analyzing condition-rich experimental designs; RSA makes it possible to represent many experimental conditions within a single visualization and to compare them within the same analysis framework. This makes it possible to assess broad distinctions (e.g., attention vs. passive) alongside finer-grained contrasts (e.g., left vs. right spatial attention) in a unified representational space.

Most previous RSA studies have focused on comparing perceptual processing as a function of the external stimuli (e.g., different visual objects, the timing of audiovisual inputs, etc.), while comparatively little work has examined how representational structure changes with internal states such as attention. Moreover, applying RSA to EEG using diverse feature types–each potentially reflecting different mechanisms of attentional processing–remains in its early stages.

An et al. (2023) examined representational patterns associated with spatial vs non-spatial auditory attention and showed that ERP and alpha-band power exhibited distinct spatiotemporal dynamics during cue presentation. However, they did not investigate how these features evolve during the subsequent period of active attention, nor how each feature uniquely contributes across the task. Building on this, we applied RSA to a condition-rich EEG dataset from the same auditory attention task explored in An et al. (2023). Our goal was to probe the spatiotemporal dynamics with which attentional control uses different dimensions of the acoustic inputs. We measured representations of auditory attention in evoked and induced neural activity, describing their changes over time and their scalp distributions. We further identified key dynamics unique to evoked and induced neural activity, advancing the understanding of how different neural signals contribute to selective auditory attention.

## 2. Methods

### 2.1. Data Collection

#### 2.1.1. Participants

Our dataset included thirty participants (16 men; 14 women; ages 19 to 44) (An et al., 2023). All participants reported normal hearing and no history of neurological disorders. All experimental procedures were approved by the IRB at Boston University’s Charles River Campus, and all participants provided written informed consent.

In the task, participants had to report a target syllable (/ba/, /da/, or /ga/, each 600 ms long) among overlapping distractors selected from the same set (Figure 1). Each of the target and distractor syllables was spoken by one of four different talkers (two male and two female; all native speakers of American English) and presented at one of five possible locations (90° left, 30° left, center, 30° right, or 90° right; spatialized using a generic head-related transfer function (Gardner and Martin, 1994).

**Figure 1.**
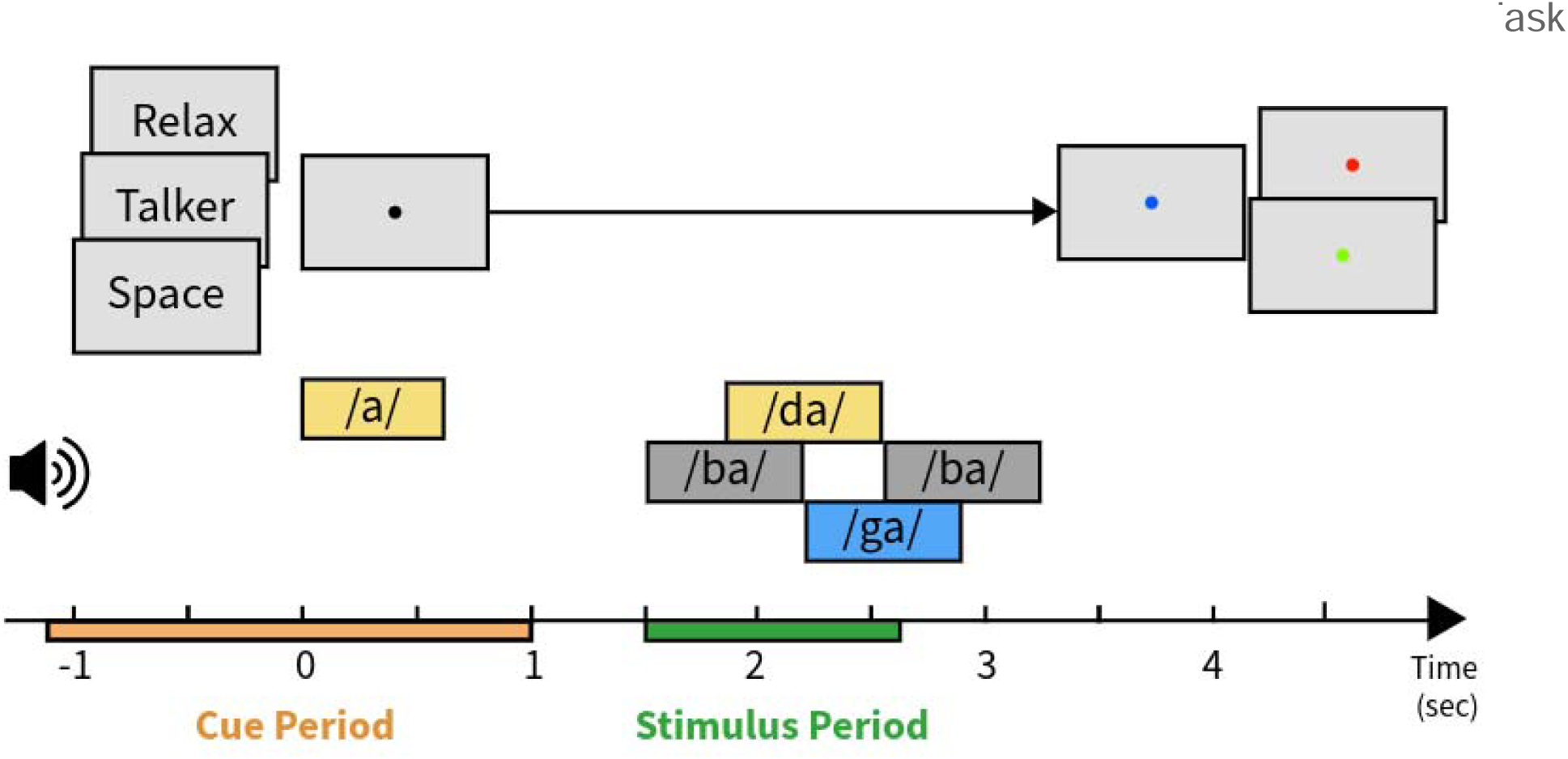
Trial structure. Each trial began with a visual task cue indicating the attention type, followed by an auditory target cue specifying either the location (spatial attention) or the talker identity (talker attention) of the upcoming target. Four overlapping auditory syllables were then played, and participants identified the syllable spoken at the target location or by the target talker. After stimulus presentation, a color change at fixation prompted the participant to make a buttonpress response. Each trial was followed by feedback (correct / incorrect). From each trial, we extract a Cue Period epoch (−1100–1000 ms from target cue onset) and a Stimulus Period epoch (−400–700 ms from target onset) for subsequent analyses.

Each trial began with a visual task cue indicating how the participant should perform the upcoming trial. “Space” indicated that spatial attention would be required; “talker” indicated that talker attention would be required; and “relax” indicated that the participant should passively listen to the stimuli but did not need to engage attention. This task cue was followed by a spoken target cue /a/ that specified the exact target. In spatial attention trials, the cue was spatialized to the exact location from which the target syllable would be presented, but spoken in a gender-neutral voice. In talker attention trials, the cue was spoken by the exact talker who would say the target syllable, but was presented at the center. In passive listening, the cue was centered and gender neutral.

After a 1000 ms preparatory period, four overlapping syllables were presented sequentially, with onsets 300ms apart. On each trial, the various syllables were randomly permuted across target and distractors; however, we always set the first and the last syllable as distractors spatialized to the center with a neutral voice, reducing primacy and recency effects and ensuring targets were masked throughout their entire presentation. The target syllable could be either the second or third syllable; the other (third or second) syllable was a *main distractor* whose spatial location and talker voice varied across conditions.

After the overlapping syllables, a color change at the fixation dot cued participants to report the target identity (1 for /ba/, 2 for /da/, 3 for /ga/), after which they received visual feedback (green for correct, red for incorrect). Participants were instructed to press any button in relax trials. The inter-trial interval was randomly set to either 1 or 1.5 seconds.

There were a total of 21 conditions: 8 “spatial”, 6 “talker”, and 7 “relax” (Table 1). Each condition comprised a specific combination of task type (spatial attention, talker attention, or passive listening), target syllable (left vs. right, female vs. male), and relationship between the target and the main distractor (e.g., how spatially close they were, whether the voices were the same or different gender). We generated 36 trials in each condition, resulting in 756 trials total. All trials were randomly shuffled, then grouped into 12 blocks for presentation.

**Table 1.** Experimental conditions. There were 21 experimental conditions defined by task type, sound location, and the talker gender of the target and distractors. In the passive listening trials, conditions were designed to match their counterparts in the attention conditions.

| # | Task | Location of Target | Location of Main Distractor | Gender of Target and Main Distractor |
| --- | --- | --- | --- | --- |
| 1 | Spatial attention | 90° Left | 30° Left | Different |
| 2 |  |  |  | Same |
| 3 |  |  | 90° Right | Different |
| 4 |  |  |  | Same |
| 5 |  | 90° Right | 30° Right | Different |
| 6 |  |  |  | Same |
| 7 |  |  | 90° Left | Different |
| 8 |  |  |  | Same |
| # | Task | Talker Gender of Target | Talker Gender of Main Distractor | Location of Target and Main Distractor |
| 9 | Talker attention | Female | Male | Co-located |
| 10 |  |  |  | Different location on the same side |
| 11 |  |  |  | Different sides |
| 12 |  | Male | Female | Co-located |
| 13 |  |  |  | Different location on the same side |
| 14 |  |  |  | Different sides |
| # | Task | Location of 2nd and 3rd syllable |  | Talker Gender of 2nd and 3rd syllable |
| 15 | Passive listening | Different locations on the left side |  | Different |
| 16 |  |  |  | Same |
| 17 |  | Different sides |  | Different |
| 18 |  |  |  | Same |
| 19 |  | Different locations on the right side |  | Different |
| 20 |  |  |  | Same |
| 21 |  | Co-located |  | Different |

Before the main experiment, participants practiced the attention tasks on a laptop. Participants repeated test runs of 21 trials (i.e., one trial for each condition) until they reached 75% accuracy. All audio was presented through a pair of insert earphones (ER1, Etymotic Research) in a sound-attenuated booth. Visual stimuli were presented on an LCD monitor within the booth.

#### 2.1.3. EEG acquisition and preprocessing

EEG was recorded using a 64-channel Biosemi ActiveTwo system, following the standard 10-20 electrode placement. All signals were sampled at 2048 Hz. After recording, EEG signals were first re-referenced to the average of two mastoid channels, then bandpass filtered between 0.5 Hz and 50 Hz using a zero-phase filter with a Hamming window, and downsampled to 512 Hz. Bad channels were rejected based on a predefined rule: channels whose maximum absolute amplitude exceeded 400µV and whose standard deviation was larger than three times the average of all channels. Rejected channels were interpolated using a spherical spline method. This yielded a minimally processed dataset.

In order to identify and reject oculomotor artifacts, we applied independent component analysis (ICA) and the automated artifact-labeling tool ICLabel. To improve the quality of ICA decomposition and the automatic labeling (Pion-Tonachini et al., 2019), we computed the ICA weights using the infomax algorithm on a temporary copy of the data that was bandpass filtered between 1 Hz and 30 Hz. Interpolated bad channels were excluded from ICA because interpolation reduces the effective rank of the data. The resulting independent components were ordered according to explained variance, and automatically labeled by ICLabel, a deep-learning-based artifact labeling algorithm (Pion-Tonachini et al., 2019). Components that were within the top 15 variance-explaning components and classified as ‘Eye’ with more than 85% confidence were identified for removal. Once computed, these ICA weights were applied back to the minimally processed EEG data; this corrected data was used for all subsequent analyses.

Time-varying power spectra for each trial were estimated using Morlet wavelets from 2 to 50 Hz (divided into 96 logarithmically spaced frequencies with a scale spacing of 0.05), with the number of cycles dynamically adjusted relative to frequency. Within each trial, power was averaged within five frequency bands: delta (2–4 Hz), theta (4–8 Hz), alpha (8–14 Hz), beta (14–20 Hz), and gamma (20–50 Hz). Two epochs were extracted from each trial: “Cue Period” epochs incorporate the task and target cue presentation (from −1100 ms to 1000 ms relative to target cue onset) and “Stimulus Period” epochs incorporate the presentation of the target syllable (from −400 ms to 700 ms relative to its onset). Because we were particularly interested in the dynamics of neural control as listeners prepared to deploy selective attention, the cue epoch is approximately twice as long as the stimulus epoch. Both epochs were baseline corrected using a common baseline window: −500 ms to −200 ms before the visual task cue.

While we investigated representations of auditory selective attention in all five frequency bands, the results below present only representations within the alpha band (8–14 Hz) and the broadband scalp voltage, as their contributions to attention representation were by far the strongest. Results in other frequency bands are presented in the supplementary materials.

Note that this preprocessing pipeline worked in a semi-automated manner for reproducibility. All preprocessing was done in Python 3.12.9 with MNE-Python 1.9.0. Raw and preprocessed EEG data are available at https://doi.org/10.1184/R1/c.8200769, and code for preprocessing and analysis is available at https://github.com/LiMN-CMU/auditory-attention-RSA.

### 2.2. Representational Similarity Analysis (RSA)

#### 2.2.1. Decoding-based Representational Dissimilarity Matrices (RDMs)

From the preprocessing, we obtained six time-varying EEG features: broadband EEG scalp voltage and power in five frequency bands (delta, theta, alpha, beta, and gamma). For each EEG feature, the dataset comprised participants, conditions, trials per condition, recorded across channels over time points (which varied depending on the epoch type).

To gain insight into the dynamics of auditory attentional control, we computed representational dissimilarity matrices (RDMs) for each participant, each EEG feature, and each time point within the epochs (see Figure 2 for an overview of the analysis pipeline). We operationalized dissimilarity as the accuracy with which a linear support vector machine (SVM) was able to classify trials. That is, we selected one participant, EEG feature, and time point, and iterated over each possible pair from the set of conditions (210 condition pairs). Employing leave-one-trial-out cross-validation (36^2^ = 1296 folds), we trained the SVM to discriminate between the two conditions. The classification accuracy was averaged across all folds to yield a dissimilarity value for that specific condition pair, participant, feature, and time point. This process was repeated for each of the 210 condition pairs, each of the 30 participants, each of the 6 EEG features, and each of the time points. Statistical analyses are performed at the level of individual subject RDMs; visualizations show grand average RDMs created by combining across all subjects.

**Figure 2.**
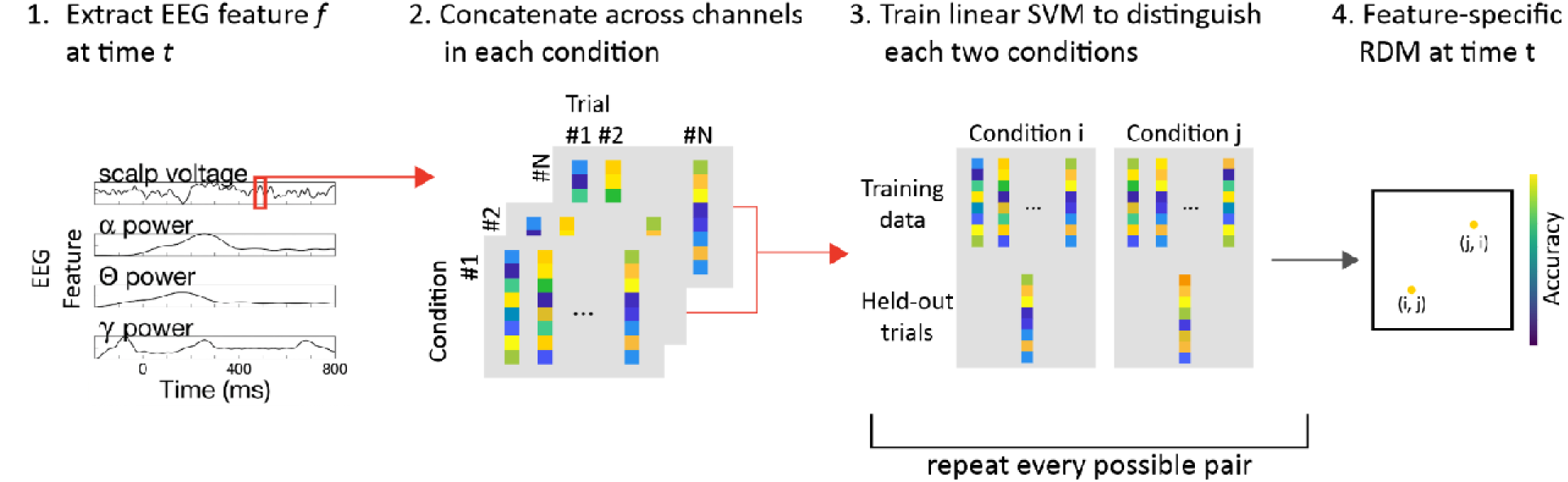
Analysis pipeline of representational similarity analysis (RSA). For each time point t, we (1) extract an EEG feature f (i.e. scalp voltage or alpha-band power) from all channels for every trial and condition, (2) concatenate the feature values across channels to form a multivariate sensor-space pattern for each trial, (3) train a linear SVM to discriminate each pair of conditions using these multivariate patterns, and (4) use pairwise decoding performance to populate the representational dissimilarity matrix (RDM) at time t. Repeating this procedure across time points yields time-resolved RDMs that summarize the separability of all condition pairs over time.

SVMs were fit separately for each time point and feature to avoid information leakage across time or features. The SVM regularization parameter of 1 was selected a priori, but lambda values from to 1 at representative time points (500ms after the target cue for the Cue Period, 400ms after the target onset for the Stimulus Period) yielded highly similar RDMs (mean pairwise correlations ≥ 0.75 during the Cue Period and ≥ 0.89 during the Stimulus Period), confirming that our results are robust to the specific choice of regularization value.

An example grand average RDM is shown in Figure 3a. Each cell in the matrix represents the dissimilarity between a pair of conditions, operationalized as the mean accuracy with which a classifier is able to distinguish between the two. Higher decoding accuracy indicates that the neural activity measured under those two conditions is more discriminable, which implies that the two conditions have distinct neural representations.

**Figure 3.**
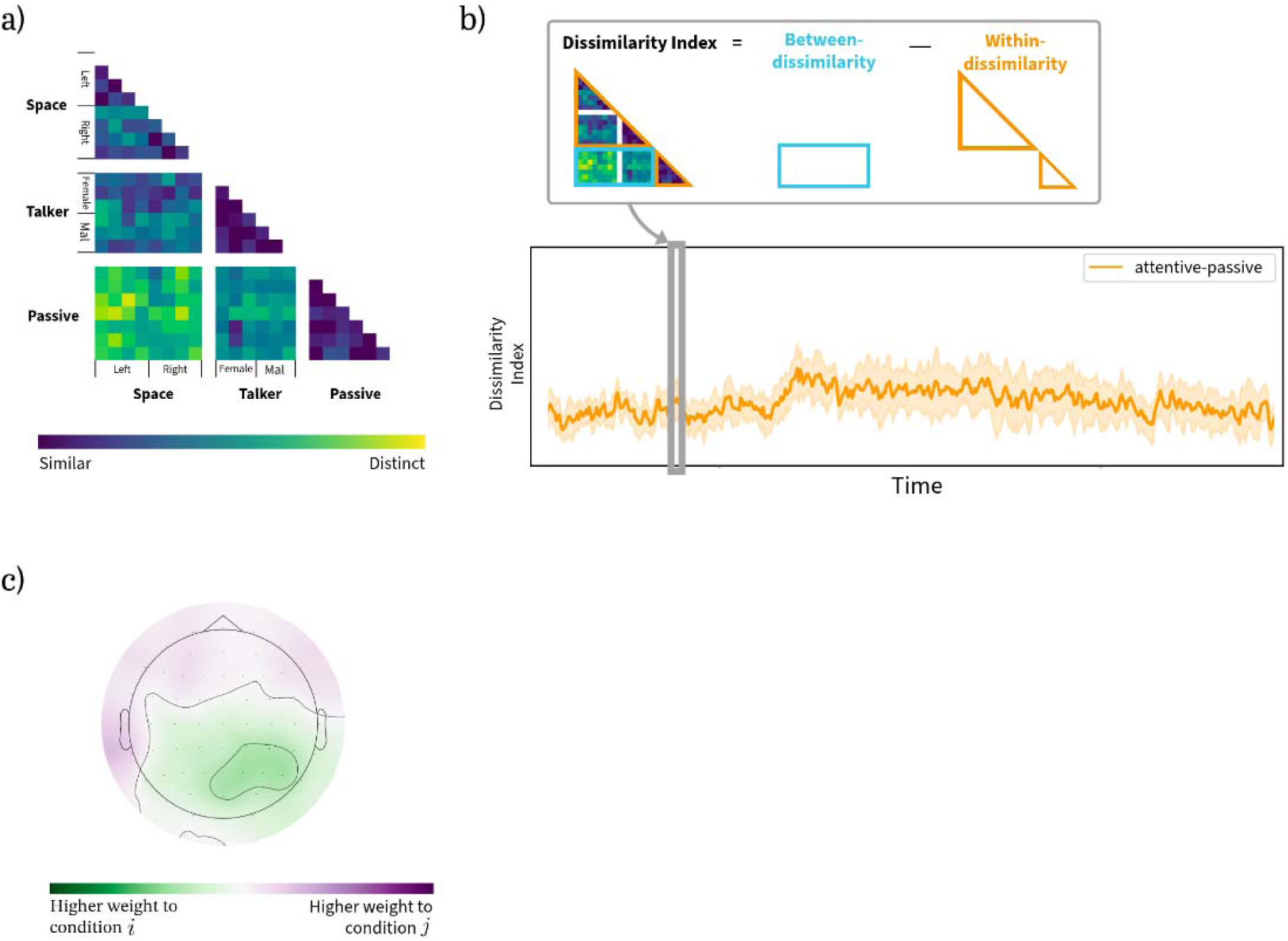
Example results from representational similarity analysis (RSA). (a) Representational dissimilarity matrix (RDM). Each cell indicates the dissimilarity (classifier accuracy) between one pair of conditions. Conditions are given in the same order as in Table 1, and are grouped for visualization by their attention task type (spatial attention, talker attention, passive listening) and primary target feature. (b) Time-resolved dissimilarity index. The top schematic illustrates the derivation of dissimilarity index (Cichy et al., 2014) which quantifies the difference between within-condition and between-condition dissimilarity by subtracting the average within-condition decoding accuracy from the between-condition average. This index reflects the similarity of attention-related representations over time. The bottom panel shows an example time course of this index. (c) Spatial importance map. Haufe-transformed classifier channel weights indicate how strongly each channel contributes to decoding. Positive values indicate that a channel contributes more strongly to condition, negative values to condition, and values near zero suggest the channel is not informative for distinguishing two conditions.

To visualize representational changes across the course of a trial, we created three dissimilarity indices quantifying how strongly different aspects of attention are represented (after Cichy et al., 2014; Figure 3b). Each dissimilarity index is computed by first averaging classifier accuracy across all cells where the compared conditions *differ* along the dimension of interest. From this value, we subtract the mean classifier accuracy across all cells where the compared conditions *share* that dimension. We computed dissimilarity indices for three contrasts: attentive listening (conditions 1-14) versus passive listening (conditions 15-21); spatial attention (conditions 1-8) versus talker attention (conditions 9-14); and attention to left hemifield targets (conditions 1-4) versus to right hemifield targets (conditions 5-8). For example, for attentive versus passive listening, we first average decoding accuracy across all pairs of conditions in which one condition involves attentive listening and the other involves passive listening, and then subtract the mean accuracy across pairs in which both conditions are attentive or both are passive. In the Results, time courses of these dissimilarity indices are presented alongside RDMs from key time points. For visualization, we plotted 95% within-subject confidence intervals computed using Cousineau–Morey method (standard error of the normalized data with Morey’s correction; Cousineau, 2005; Morey, 2008).

Dissimilarity indices were tested for significant differences from zero using one-dimensional cluster-based permutation tests (Maris & Oostenveld, 2007). At each time point, we computed the *t*-statistic against zero; adjacent time points with *t*-value > 2.756 (corresponding to an uncorrected two-tailed *p* < .01 for df = 29) were grouped into clusters; cluster-level statistics were defined as the sum of *t*-values within that cluster. A null distribution of cluster statistics was created using 1,000 random permutations of condition labels, and *p*-values were computed by comparing the observed clusters against this distribution. Statistical significance was determined at *p* < 0.05 with Holm-Bonferroni correction for multiple comparisons (Holm, 1979).

#### 2.2.2. Spatial importance maps

In order to understand which EEG channels carry the strongest neural representations, we projected the classifier’s weights back into channel space to visualize how these spatial patterns change over the course of the task and differ across condition contrasts. Since classifiers were trained independently at every time point, consistent weights across time may reflect persistent underlying processes.

As described above, we trained SVMs to classify each pair of conditions for each subject, EEG feature, and time point using leave-one-trial-out cross-validation. For the current analysis, we averaged the SVM weight vectors across cross-validation folds, yielding one 64-channel weight vector for each condition pair. We then applied the Haufe transformation (Haufe et al., 2014) to convert these averaged decoding weights into encoding patterns that are more directly interpretable in channel space, as raw classifier weights can be influenced by noise correlations and feature scaling rather than reflecting signal strength at each channel. We again constructed three contrasts: attentive listening versus passive listening, spatial attention versus talker attention, and attention to left versus right spatial hemifield. For each contrast of interest, we averaged the Haufe-transformed weights from all relevant condition pairs in that contrast. For example, for the attentive-passive contrast, we averaged weights from all classifiers distinguishing an attentive condition from a passive condition. We then averaged these contrast-specific weight maps across participants to create grand-average spatial importance maps for each time point.

An example spatial map is shown in Figure 3c. Positive values (green) indicate channels more strongly associated with one condition, whereas negative values (purple) indicate channels more strongly associated with the other. Values near zero indicate minimal contribution to discriminating the two conditions. Thus, the sign indicates which side of the contrast a channel is more strongly aligned with, whereas the magnitude indicates how strongly that channel contributes to the discriminative spatial pattern. These maps characterize channels in terms of their contribution to discriminating different conditions, unlike conventional EEG topographies, which show the amplitude of an EEG feature at each channel.

#### 2.2.3. Unique contribution analysis

Alpha power has been associated with selective attention in many studies (e.g., Deng et al., 2020; Kerlin et al., 2010; Wöstmann et al., 2021) and represented attention most strongly in our data (see Results). To assess whether the representation in other EEG features was complementary or redundant to that in alpha power, we compared RDMs derived from dual-feature decoding (using both alpha power and one other feature) to those derived using each feature independently. Alpha power was concatenated with one additional feature (EEG scalp voltage, delta-, theta-, beta-, or gamma-band power) along the channel axis, resulting in a 128-channel weight vector. A stronger regularization parameter of 0.01 was used to account for the doubled number of inputs. The rest of the parameters and the training procedure remained the same.

To evaluate how evoked and induced EEG activity each contribute to the representation of attention, we quantified how much decoding accuracy with dual-feature decoding exceeded accuracy with a single feature (Di Liberto et al., 2018). For example, the unique contribution of alpha-band power (induced) relative to broadband voltage (evoked) was computed by subtracting the RDM based on broadband voltage alone from the RDM derived from both features together. Representational structure present in the dual-feature but not the single-feature RDM can thus be attributed to the unique contribution of alpha-band oscillations. The converse was done to estimate the unique contribution of evoked activity relative to induced alpha-band activity. As above, we computed dissimilarity indices to summarize these RDMs over time and identified windows of statistically significant contributions using cluster-based permutation testing.

The main manuscript presents the unique contribution analysis of alpha-band power and broadband scalp voltage; analysis of alpha compared to other frequency bands is presented in the supplemental materials.

## 3. Results

### 3.1. Behavior

All participants achieved high accuracy in recognizing the target syllable (Mean=0.88, SD=0.07 in spatial attention, Mean=0.94, SD=0.05 in talker attention). Thus, we included all trials—both correct and incorrect—in subsequent analyses. Detailed information can be found in An et al. (2023).

### 3.2. Representational Dissimilarity of Attention

To probe the spatiotemporal dynamics of attention during a challenging auditory attention task, we measured two neural features (evoked broadband scalp voltage and induced alpha-band power) and computed timepoint by timepoint representational dissimilarity in two epochs: the Cue Period epoch spanning the task cue, target cue, and the time leading up to stimulus presentation, and Stimulus Period epoch surrounding the presentation of the target syllable (see Methods).

#### 3.2.1. Cue Period

We first examined the Cue Period during which listeners prepared to deploy selective attention. Neural representations reflected in evoked broadband scalp voltage and in induced alpha-band power showed distinct time dynamics. Scalp voltage (Figure 4a–c) exhibited a rapid increase immediately after the target cue, followed by a decline over time, whereas alpha-band power showed a more gradual increase that persisted through the Cue Period.

**Figure 4.**
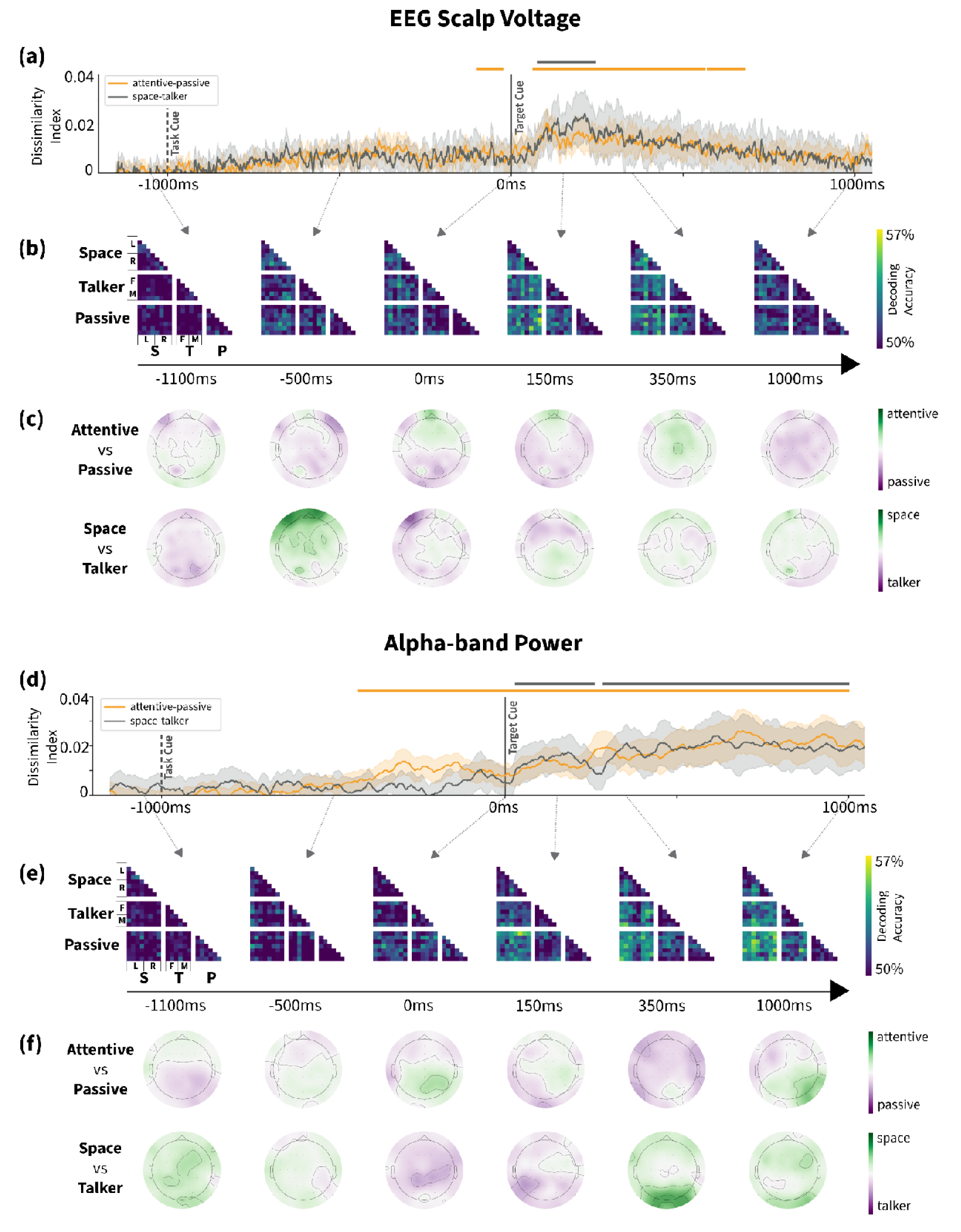
Representational dissimilarity during Cue Period in broadband scalp voltage (top) and alpha-band power (bottom). (a,d) Dissimilarity indices over time for the contrast between attentive and passive listening (orange) and between spatial and talker attention (gray). Error bands are 95% within-subject, standard error-based confidence intervals of the mean. Onsets of the task cue (−1000 ms, dashed line) and target cue (0 ms, solid line) are indicated. Horizontal lines denote temporal clusters of statistical significance. (b,e) Representational dissimilarity matrices (RDMs) at representative time points throughout the Cue Period. (c,f) Spatial importance maps illustrating channel-level contributions to SVM classification at each representative time point.

In broadband scalp voltage, there was no representation of task before the onset of any trial-specific information, 1100 ms before target cue onset. The attentive-passive dissimilarity index became significantly positive approximately 400 ms after task cue onset and remained elevated for nearly the entire epoch. The space-talker dissimilarity index hovered near zero until the target cue occurred. Both dissimilarity indices peaked around 100–200 ms after target cue onset and then decreased over time. Note that clusters shorter than 30 ms were removed to reduce the influence of short-term noise. Representational dissimilarity matrices (RDMs) at 150ms and 350ms after target cue show clear structure: specifically, spatial attention representations are most distinct from other conditions. In the same period, spatial importance maps displayed higher weights in the frontocentral area for the space-talker attention contrast.

In alpha-band power (Figure 4d–f), the attentive-passive dissimilarity index increased following the task cue, becoming significantly greater than zero at approximately 500 ms after task cue and remaining positive until the end of the Cue Period. Shortly after target cue onset, the space-talker index also increased, becoming significantly greater than zero at around 50 ms and persisting until the end of the Cue Period. The RDM at 1000 ms after target cue shows clear separation for both contrasts (attentive-passive and spatial-talker attention). In the corresponding spatial importance maps, parietal-occipital channels carry higher weights for both contrasts 1000 ms after target cue.

Lower frequency bands (delta and theta) followed a pattern similar to scalp voltage, with peaks around 100–200 ms and a rapid attenuation. In contrast, decoding accuracies based on beta-and gamma-band power remained generally low throughout the period (Supplementary Figure 1).

Because we expected the anticipated spatial location of an upcoming target to be represented in the distribution of alpha-band oscillations (as in Deng et al., 2020; Gutteling et al., 2022), we examined the contrast between spatial attention to a left hemifield target and to a right hemifield target during the Cue Period (Figure 5). The left-right dissimilarity index is significantly above zero 350–800 ms after target cue onset; the RDMs at these times also reflect the encoding of the direction of attentional focus. The spatial importance maps identified lateralized activity in the parietal region at 350 ms and 500 ms after target cue onset.

**Figure 5.**
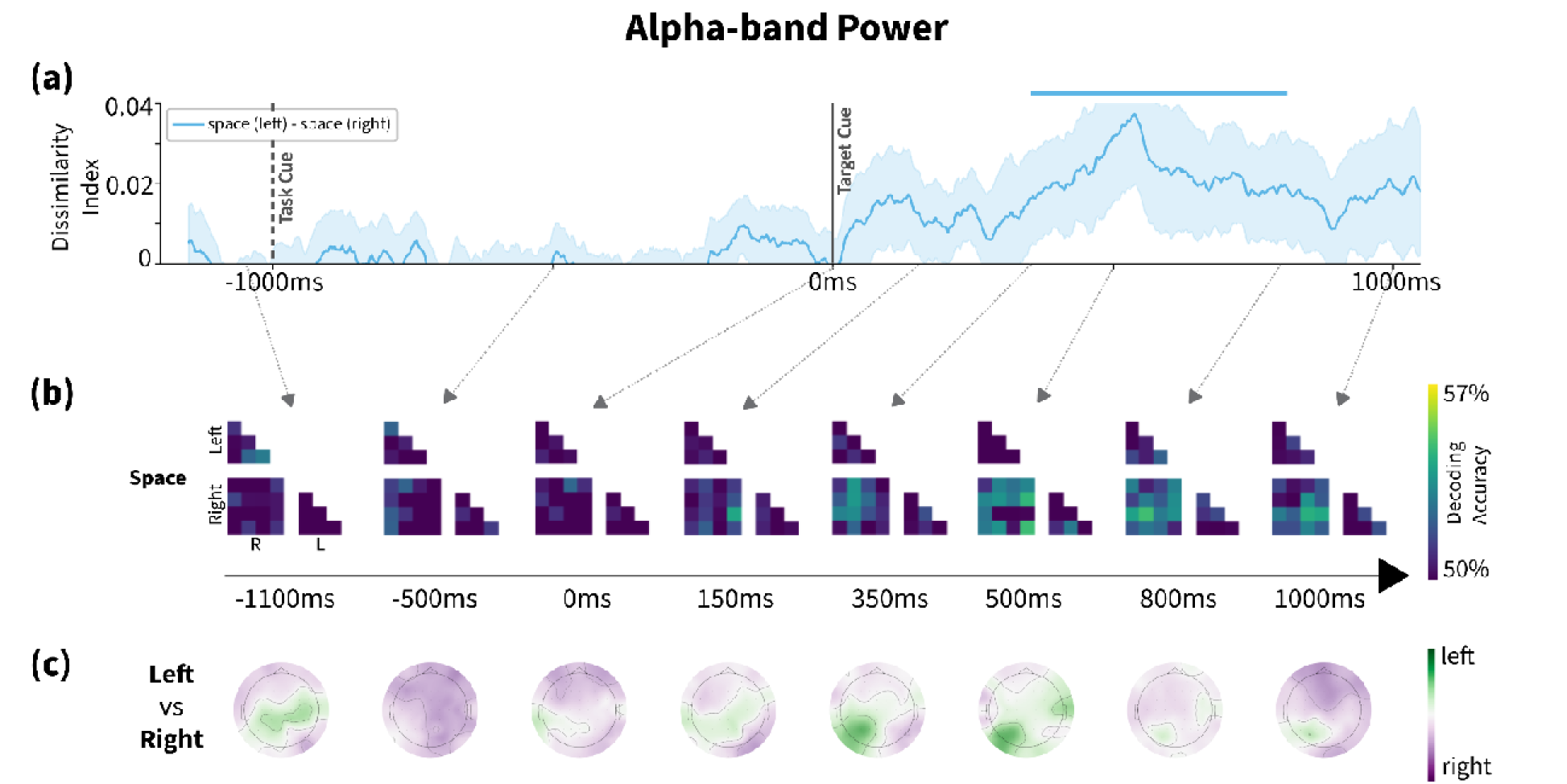
Representational dissimilarity in alpha power between left and right spatial attention during Cue Period. (a) Dissimilarity indices over time for spatial attention trials with left versus right hemifield targets. Error bands are 95% within-subject, standard error-based confidence intervals of the mean. Onsets of the task cue (−1000 ms, dashed line) and target cue (0 ms, solid line) are indicated. Horizontal lines denote temporal clusters of statistical significance, (b) Representational dissimilarity matrices (RDMs) at representative time points throughout the Cue Period for spatial attention conditions. (c) Spatial importance maps illustrating channel-level contributions to SVM classification at those representative time points.

#### 3.2.2. Stimulus Period

Our second epoch of interest encompassed the target syllable and its immediate neighbors, capturing neural activity during active selective attention. Overall, neural representations of attention remained significantly above zero throughout this period in both evoked broadband scalp voltage and induced alpha-band power, but their temporal dynamics differed.

In scalp voltage (Figure 6a–c), the attentive-passive dissimilarity index was positive throughout this period, but increased markedly shortly after the onset of the target syllable, remaining elevated from approximately 300 ms after syllable onset until the end of the epoch. The spatial importance maps showed a consistent central-posterior distribution until 200ms, but this pattern shifted toward posterior sensors around 400 ms. The RDMs also directly confirmed this, with the strongest dissimilarity values emerging late in the Stimulus Period. Space-talker dissimilarity, however, remained largely non-significant except for a transient period 300–400 ms after target onset, and its peak was much lower than for other contrasts.

**Figure 6.**
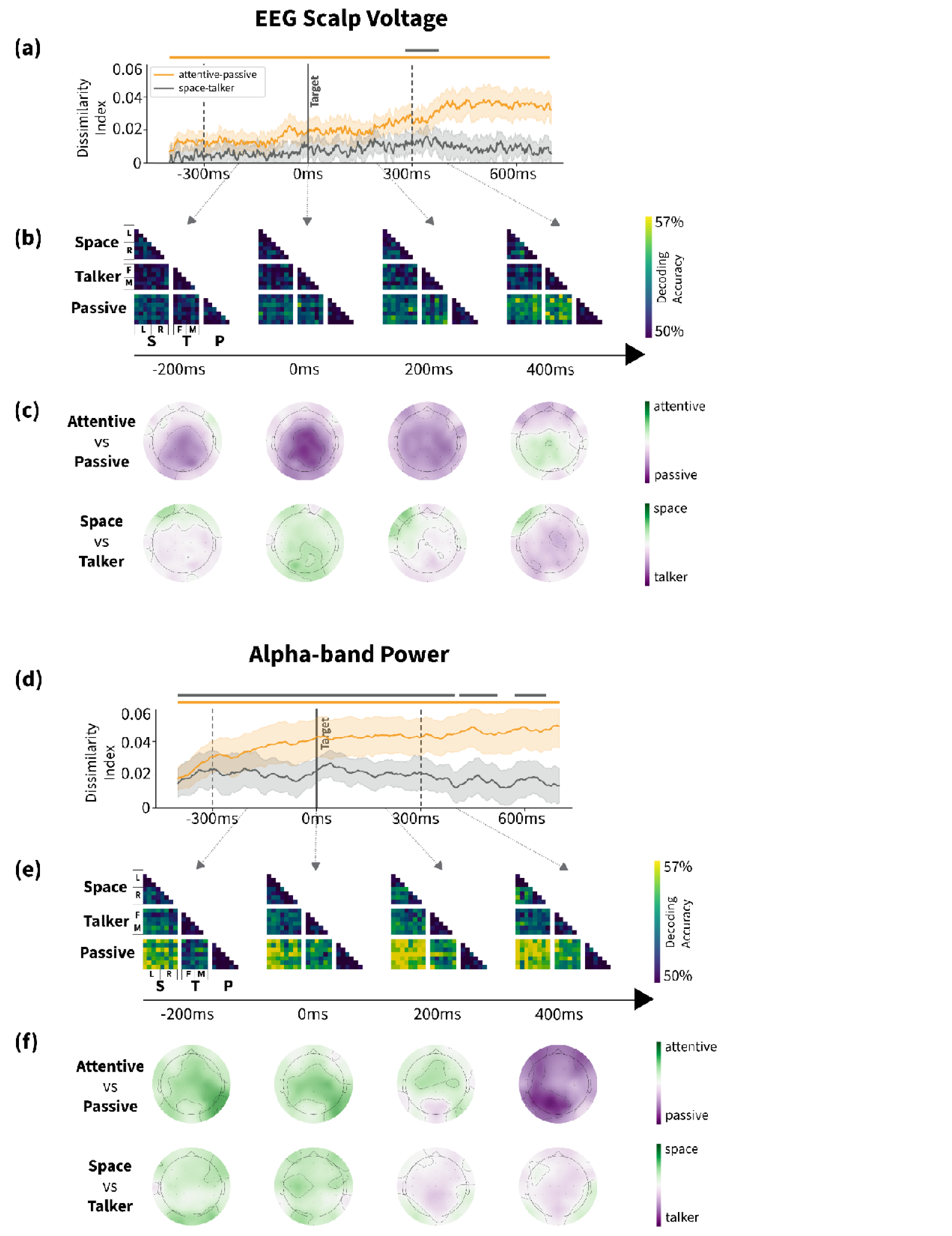
Representational dissimilarity during Stimulus Period in broadband scalp voltage (top) and alpha-band power (bottom). (a,d) Dissimilarity indices over time for the contrast between attentive and passive listening (orange) and between spatial and talker attention (gray). Error bands are 95% within-subject, standard error-based confidence intervals of the mean. Onsets of the pre-target syllable (−300 ms, dashed line), target syllable (0 ms, solid line), and post-target syllable (300 ms, dashed line) are indicated. Horizontal lines denote temporal clusters of statistical significance. b,e) Representational dissimilarity matrices (RDMs) at representative time points throughout the Stimulus Period. (c,f) Spatial importance maps illustrating channel-level contributions to SVM classification at each representative time point.

In contrast, alpha power demonstrated high decoding accuracy throughout the entire period (Figure 6d–f), with both dissimilarity indices significantly positive. Alpha-band RDMs showed high levels of dissimilarity across conditions throughout this entire period, with little change in the degree or structure. Spatial importance maps show a broad central-posterior weighting for both contrasts until target syllable presentation, after which they diverge. Channel weights in attentive-passive contrast flip from positive to negative at 400 ms after target syllable onset, while those in the spatial-talker contrast show no clearly defined spatial pattern at either 200 and 400 ms. Similar but weaker patterns were observed in other frequency bands. (Supplementary Figure 2).

Alpha-band oscillations carry representations of target location during spatial attention (Figure 7). Left-right dissimilarity remained significantly above zero from the onset of the pre-target syllable through the end of the following one (also visible in the RDMs), especially those at 200 and 400 ms after target syllable onset. Consistent with these findings, the spatial importance maps at these times exhibited parietal lateralization similar to that observed during the Cue Period.

**Figure 7.**
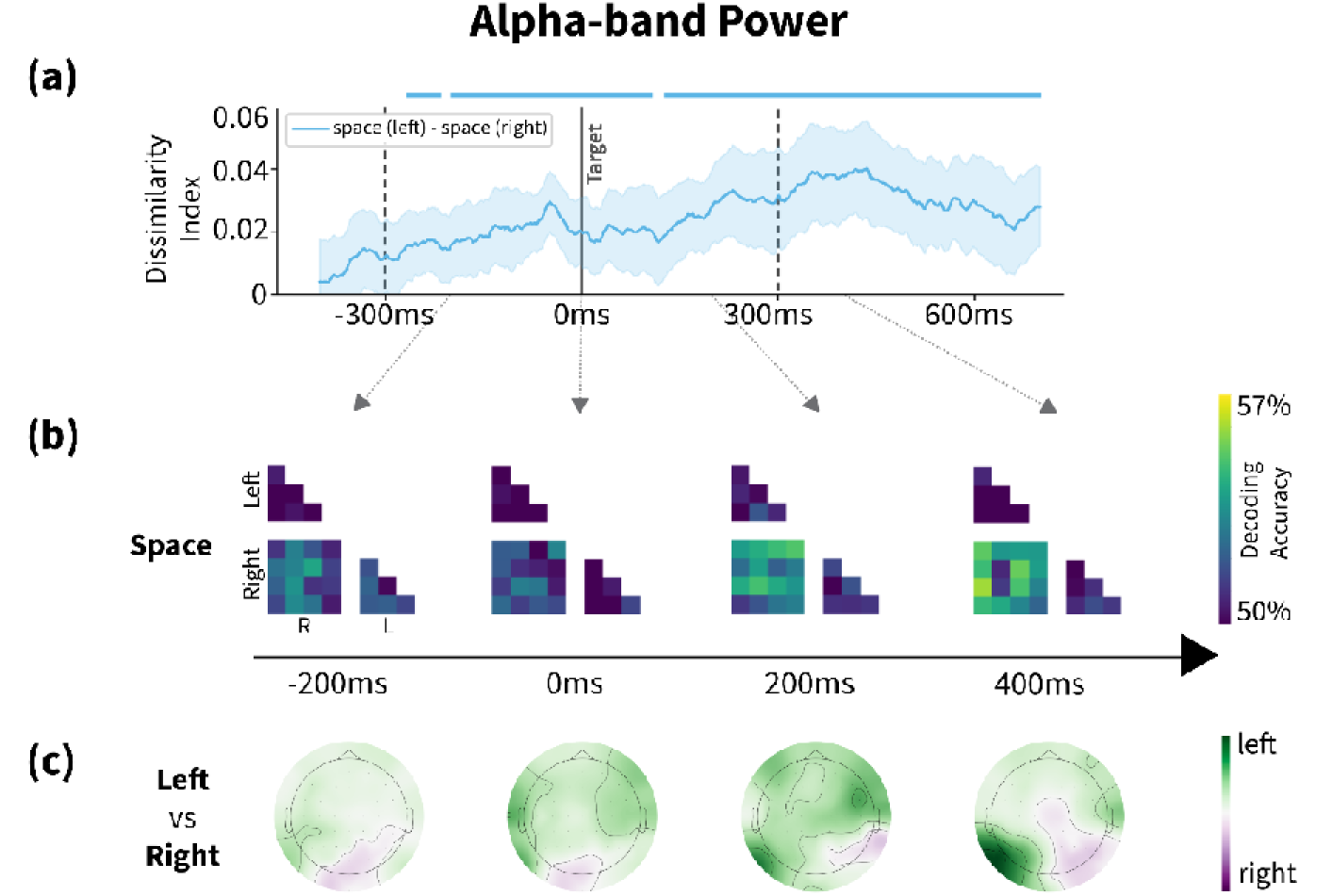
Representational dissimilarity matrices in alpha power between left and right spatial attention during Stimulus Period. (a) Dissimilarity indices over time for the contrast between spatial attention trials with a left hemifield target and with a right hemifield target. Error bands are 95% within-subject, standard error-based confidence intervals of the mean. Onsets of the pre-target syllable (−300 ms, dashed line), target syllable (0 ms, solid line), and post-target syllable (300 ms, dashed line) are indicated. Horizontal lines denote temporal clusters of statistical significance. (b) Representational dissimilarity matrices (RDMs) at representative time points throughout the Cue Period for spatial attention conditions. (c) Spatial importance maps illustrating channel-level contributions to SVM classification at those representative time points.

### 3.3. Unique Contribution of EEG Features

To measure whether broadband scalp voltage and EEG alpha power are carrying redundant or complementary information, we estimated timepoint by timepoint RDMs from both features combined (Supplementary Figures 3 and 5), then subtracted single-feature RDMs at the corresponding timepoints. That is, to estimate the information that is represented uniquely in alpha oscillations, we subtracted the broadband RDMs from the dual-feature (broadband + alpha) RDMs. The converse operation yielded RDMs describing the information contributed uniquely by broadband activity.

#### 3.3.1. Cue Period

In the Cue Period, evoked broadband scalp voltage and induced alpha power exhibited complementary time courses. The unique contribution of scalp voltage (Figure 8a) was short-lived, with brief periods of significant dissimilarity within 500 ms following the onset of target cue. In contrast, the unique contribution of alpha power (Figure 8b) developed more slowly but was sustained, becoming significantly positive around 200 ms after target cue and remaining robust throughout the period.

**Figure 8.**
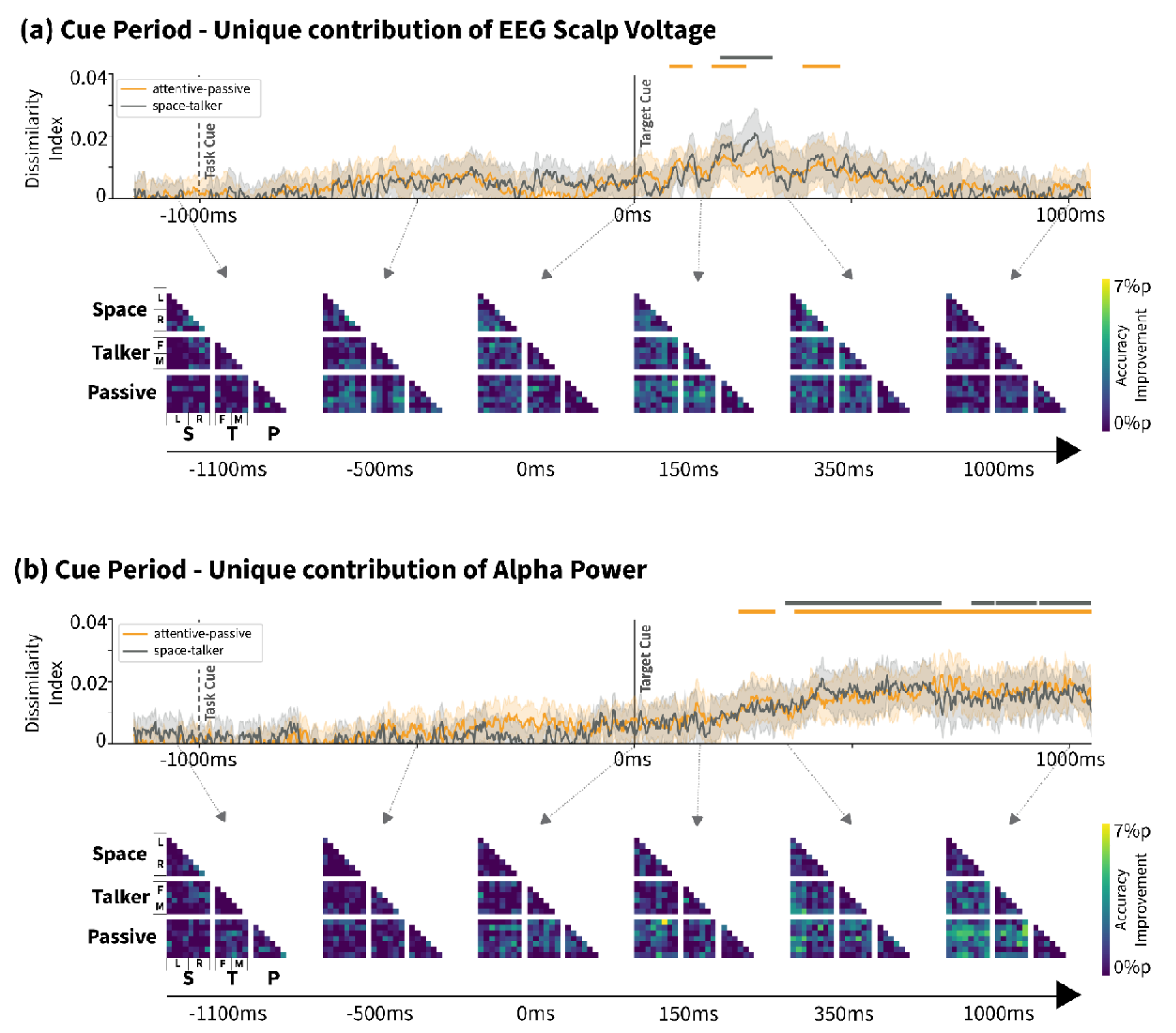
Unique contribution of (a) broadband scalp voltage and (b) alpha power during Cue Period. Upper panel: Dissimilarity indices over time for the contrast between attention and passive (orange) and between spatial and talker attention (gray), summarizing the information carried in one EEG measurement above that carried by the other. Error bands are 95% within-subject, standard error-based confidence intervals of the mean. Onsets of the task cue (−1000 ms, dashed line) and target cue (0 ms, solid line) are indicated. Horizontal lines denote temporal clusters of statistical significance. Lower panel: Unique contribution RDMs at representative time points throughout the Cue Period.

Similar analyses compared alpha power representations to those in other frequency bands and found that improvements were mostly driven by alpha, with little contribution from the others except theta, which displayed a brief peak immediately following the target cue (Supplementary Figures 4).

#### 3.3.2. Stimulus Period

During stimulus presentation, the contribution of EEG scalp voltage remained low until –300 ms after the target syllable onset (Figure 9a). After 300 ms, the attentive-passive listening dissimilarity index increased and reached statistical significance. The spatial-talker attention contrast was not significant at any time point.

**Figure 9.**
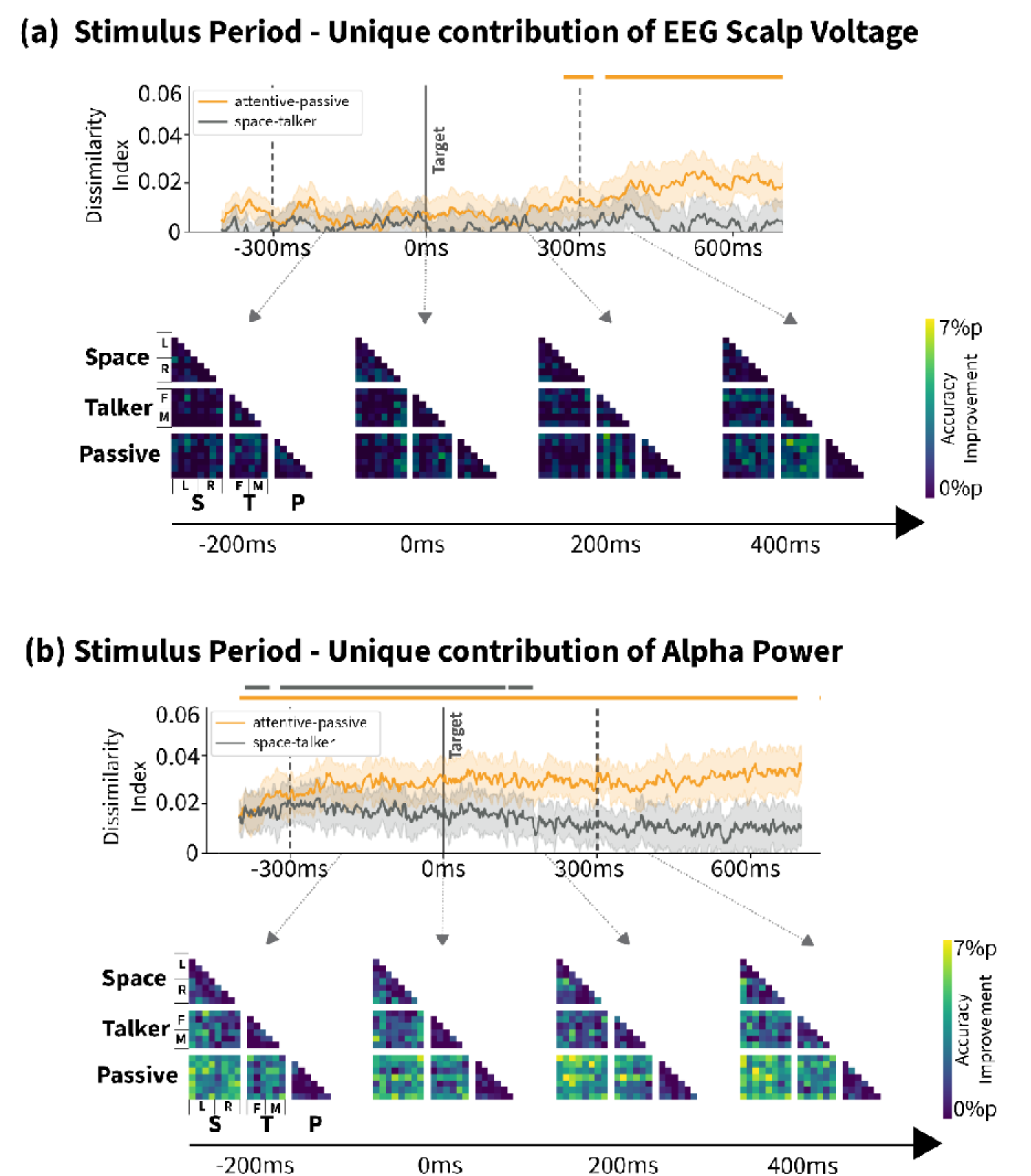
Unique contribution of (a) broadband scalp voltage and (b) alpha power during Stimulus Period. Upper panel: Dissimilarity indices over time for the contrast between attention and passive (orange) and between spatial and talker attention (gray), summarizing the information carried in one EEG measurement above that carried by the other. Error bands are 95% within-subject, standard error-based confidence intervals of the mean. Onsets of the pre-target syllable (−300 ms, dashed line), target syllable (0 ms, solid line), and post-target syllable (300 ms, dashed line) are indicated. Horizontal lines denote temporal clusters of statistical significance. Lower panel: Unique contribution RDMs at representative time points throughout the Stimulus Period.

Alpha power remained the dominant contributor to decoding throughout the period (Figure 9b). Attentive-passive dissimilarity was significant across the entire period, while the spatial-talker attention contrast was significant for approximately 600 ms surrounding the target onset. Other frequency bands provide little to no additional information beyond that carried in alpha (Supplementary Figures 6).

## 4. Discussion

We demonstrated that representational similarity analysis (RSA) can effectively capture the dynamics of both evoked and induced EEG activity during an auditory attention task. Broadband scalp voltage revealed two stimulus□evoked representational patterns: one emerging shortly after the target identity cue, and another following the onset of the target syllable. In contrast, representations in induced alpha-band activity developed gradually and persisted throughout the trial. Unique contribution analysis revealed that these two types of neural activity carried complementary representations, each encoding information beyond that carried by the other.

By applying RSA to internal cognitive states, this study examines the neural representations of task goals and attentional control. We minimized effects of acoustic difference across conditions by carefully matching the auditory stimuli while varying the cue dimension that would identify the target. This design allowed us to isolate changes in representation that were attributable to internal task goals and differences in how attention was deployed.

### Neural representations in broadband scalp voltage show patterns similar to auditory event-related potentials

Transient representational differences between attention and passive listening conditions were apparent in broadband voltage shortly after the auditory cue (Figures 4a–c). Previous work has shown that attention modulates auditory evoked responses within 100-300 ms of sound onset (Choi et al., 2013; Hillyard et al., 1973; Nguyen et al., 2024, Näätänen & Picton, 1987), which is the time window in which we observed this effect. In the sensory gain control framework (Hillyard et al., 1973, Hillyard et al., 1998), attention selectively modulates sensory responses in auditory cortex such that evoked responses to a particular sound are relatively larger when that sound is attended compared to when it is ignored. Our spatial importance maps showed relatively higher classifier weights in frontocentral channels at 150 ms and 350 ms post-cue when comparing attentive to passive listening. These channels typically show attentional modulation of auditory evoked responses in traditional EEG experiments (Hillyard et al., 1973; Näätänen & Picton, 1987).

Throughout stimulus presentation, the attentive-passive dissimilarity was reliably significant, but dissimilarity increased from approximately 300 ms after target onset (Figure 6a). This time window is consistent with the evoked P300 target-detection response, which arises in centro-parietal channels after a task□relevant target is detected (Donchin & Coles, 1988). The P300 may reflect working memory updates when important or task-relevant information is perceived (Polich, 2007; Watter et al., 2001; Guerrero et al., 2023). Consistent with this interpretation, spatial importance maps show high central-parietal weights in attentive-passive contrasts at this time. However, other processes such as motor planning or short-term memory maintenance may also contribute to this response (Verleger et al., 2005; Ruchkin et al., 1992).

Representational differences between spatial and talker attention were only reliably significant shortly after the target identity cue (around 100–300 ms, see Figure 4a). However, the target identity cues were not as carefully matched as other stimulus parameters. In spatial attention trials, this cue was lateralized to the target location with a neutral voice; in talker attention and passive listening trials, this cue was presented at the center with the same voice as the target syllable. ERPs elicited by lateralized sounds are larger than for medial sounds, even under passive listening (Näätänen & Picton 1987). This decoding success may thus reflect physical properties of auditory stimuli (i.e., source laterality) rather than (or in addition to) differences in the internal states of participants. During stimulus presentations where both target and distractor were lateralized, the space-talker dissimilarity was not reliably significant. We interpret this pattern as indicating that the evoked activity represents attentional engagement but does not represent the task-relevant feature to a degree that can be decoded from scalp voltage.

### Neural representations in alpha-band power showed consistent patterns from preparatory period to stimulus presentation

Representational differences extracted from alpha power emerged gradually after the target cue and persisted throughout the trial. RDM patterns were strong, particularly during the Stimulus Period when attentional engagement was likely highest. Representations in alpha power were more distinctive than those in any other EEG feature, consistent with many prior findings linking alpha oscillations to attention (e.g., Deng et al., 2020; Kerlin et al., 2010; Wöstmann et al., 2021; Klimesch, 2012).

Alpha-band oscillations have been linked to a gating mechanism that shields ongoing cognitive processing from interference (Jensen & Mazaheri, 2010; Foxe & Snyder, 2011; Strauß et al., 2014). Alpha power measured at the scalp likely reflects several different neural generators that are engaged during selective attention (Deng et al., 2020; Wöstmann et al., 2017; Gutteling et al., 2022; see review in Jensen & Bonnefond, 2026). Posterior parietal and occipital alpha is strongly related to spatial attention to a specific hemifield, with alpha power increasing ipsilaterally and decreasing contralaterally to the target location, as if it is suppressing inputs at task-irrelevant locations (Deng et al., 2020; Wöstmann et al., 2019). Beyond spatial orienting, alpha power also reflects more general aspects of attentional demand and cognitive control (Wöstmann et al., 2017, 2019; Gutteling et al., 2022). For example, increases in posterior alpha accompany higher acoustic detail or disturbance in the distractor stream, consistent with a role in shielding target processing from interfering input (Wöstmann et al., 2019). Further, parieto-occipital alpha power predicts trial-wise speech intelligibility in competing sounds, linking this shielding mechanism to behavioral performance (Viswanathan et al., 2023).

Within this framework, the stable alpha-based RDM patterns from preparation through stimulus presentation suggest that alpha oscillations persistently represent attentional state. We observe robust separation of attentive versus passive conditions and spatial weighting of parietal-occipital channels, both consistent with models in which posterior alpha maintains the gain control that prioritizes certain sensory inputs for subsequent processing. In our data, alpha thus appears to index an ongoing control signal that maintains target-relevant representations over time. At the same time, our scalp-level measurements cannot isolate which neural generators are primarily responsible for the observed alpha patterns. Future work using measurements with high spatial precision or causal perturbations (e.g., parietal or frontal TMS) will be necessary to determine which nodes of the attention network actually generate the persistent alpha-band representational structure we report here.

Alpha-band activity also reflected which attentional dimension participants used: spatial versus talker identity cues were associated with different neural patterns reflected in alpha power, both during preparation and during stimulus presentation (Figure 4d and 6d). When we further separated spatial attention by target location (i.e., left vs. right hemifield), alpha-based spatial importance maps exhibited clear parietal lateralization, particularly 300–500 ms following cue onset and 300 ms after the target (Figure 5 and 7). This lateralized pattern is consistent with previous reports of alpha-band modulation during spatial attention (Deng et al., 2020; Wöstmann et al., 2017; Gutteling et al., 2022). Together, these findings suggest that distributed alpha-band dynamics carry information about both the spatial and non-spatial dimensions of auditory attention, with spatial attention exhibiting a more distinct topography.

We note that alpha power can also be modulated by factors such as overall arousal and listening effort (e.g., Obleser et al., 2012; Wöstmann et al., 2017). However, in the present study the acoustic input in the Stimulus Period was identical across tasks, the different attentional conditions were interleaved with relatively short intervals, and behavioral performance was high in both tasks. These features make it unlikely that our alpha effects can be fully explained by systematic differences in these factors between tasks.

The fact that alpha-based representations differentiate both the type of attentional cue (spatial versus talker identity) and the specific spatial focus (left versus right) shows that alpha-related processes do more than distinguish attentive from passive states; they also carry information about which acoustic dimension is guiding attention. Many studies have linked alpha increases to spatial attention in both auditory and visual tasks (Deng et al., 2020; Gutteling et al., 2022), but only a few have examined how non-spatial attention (such as talker identity) is expressed in alpha-band activity (Wöstmann et al., 2017; Viswanathan et al., 2023). Our findings therefore extend prior work on spatial alpha by demonstrating that the distributed patterns of alpha-band activity can distinguish spatial from non-spatial attentional sets such as talker-based attention.

Our results also indicate a limit to what can be decoded from EEG in this paradigm. Although talker-based attention formed a distinct representational pattern in alpha-band activity, we could not reliably discriminate between attention to female versus male talkers within the talker attention task. This may reflect both the limited spatial resolution of EEG and the distributed cortical representation of voice pitch and timbre (Burle et al., 2015; Hamilton et al., 2021). Future work that contrasts spatial selection from multiple forms of non-spatial selection (e.g., talker identity, pitch, temporal envelope, semantic category) at the representational level will be important for determining how specifically alpha-related processes code spatial versus non-spatial attention. To resolve such fine-grained talker-identity representations, modalities with higher spatial precision, such as MEG, fMRI, or intracranial EEG are likely required (for instance, see recent intracranial work on speech; van der Heijden et al., 2025).

### Information in scalp voltage and alpha-band power is unique and complementary, unlike other frequency bands

Unique contribution analysis highlighted distinct and complementary roles of broadband activity and alpha oscillations. Broadband voltage showed strong representational strength at two different time points, one near target cue onset and the other after the target syllable (Figure 8a and 9a). In contrast, representations encoded in alpha power gradually increased after the cue and remained elevated throughout the trial (Figure 8b and 9b). In particular, scalp voltage exhibited representational dissimilarity for attentive-passive contrasts, in which key information appears in a specific stimulus at a specific time. Conversely, alpha power encoded both attentive-passive and spatial-talker attention contrasts, capturing sustained task engagement and attentional state. Spatial importance maps also differed at these time points: broadband voltage showed peaks over frontocentral channels, whereas alpha power was strongest over parietal-occipital sensors. These differences remained after removing information shared between the two features.

Compared to the complementary relationship broadband scalp voltage and alpha oscillations, information in other frequency bands is largely redundant with these two signals (Supplemental figures 4–7 and 9–12). Lower frequency bands (delta, theta) showed rapid increases in discrimination accuracy during the Cue Period that faded quickly, likely reflecting ERP components within those bands. Consistent with this, theta-band spatial importance maps during the Cue Period resembled those of EEG scalp voltage. This suggests that, at least in this paradigm, much of the discriminative information in delta and theta is redundant with broadband evoked activity, rather than providing a window into any other attentional mechanisms.

Higher frequencies (beta, gamma) displayed patterns similar to those in alpha power but with considerably weaker effects. Unique contribution analysis emphasized alpha power’s dominance, revealing substantial overlap of the information conveyed by alpha and these other bands. Prior research has suggested that gamma band fluctuations contribute to speech processing, and are modulated by attention (Viswanathan et al., 2019; Synigal et al., 2020). Consistent with this, we find that high frequency features do carry information distinguishing attentive from passive listening. However, the small unique contribution of beta and gamma power (despite their visual resemblance to alpha patterns) indicates that high-frequency induced activity adds relatively little independent information about task state beyond what is already captured by alpha in this task design.

Together, these results demonstrate that RSA-based unique contribution analysis can help disentangle which EEG features truly add independent information about attentional state rather than simply re-expressing the same underlying neural dynamics in a different EEG feature.

### Limitations & Future work

We estimated dissimilarity between conditions using classifier accuracy rather than Euclidean, crossnobis, or correlation-based distance measures (Schütt et al., 2023; see Diedrichsen & Kriegeskorte, 2017 for review). We chose this approach to emphasize the decodability of neural information, making the metric interpretable: it reflects how easily information can be read out from the EEG patterns. Moreover, a multi-channel classifier yields weight maps that can highlight the sensors or spatiotemporal patterns contributing most strongly to distinguishing different states. We used a linear SVM as the decoding model because it combines moderate performance with a simple linear decision rule: it implements a weighted summation followed by a decision boundary, matching the standard linear readout assumptions often used in neuroscience and making our assumptions about downstream decoding explicit (Hung et al., 2005; Yamins et al., 2014; Diedrichsen & Kriegeskorte, 2017). Using a linear SVM also ensures that these weight maps arise from a simple linear relationship between features and classifier output and can be transformed into interpretable encoding patterns.

At the same time, using classifier accuracy introduces several limitations. Dissimilarity estimates depend on the choice of classifier, regularization, and cross□validation scheme, potentially reducing robustness and reproducibility. In our study, we mitigated this by using a simple linear model and validated regularization parameters, but the conceptual dependence remains. In addition, although Haufe□transformed weight maps make it possible to move from classifier weights to activation patterns, interpreting individual sensor values as positively or negatively associated with an outcome remains difficult, since the weights could be influenced by noise correlations and multicollinearity (Haufe et al., 2014). Future work should therefore complement accuracy□based RDMs with continuous distance measures.

Intrinsic differences in temporal resolution between scalp voltage and frequency-band features is an important limitation for our dynamic RSA. Although each RDM is computed on data from a single timepoint, the covariance between successive timepoints differs between the broadband voltage and the oscillatory power. Each estimate of power at a given time and frequency requires integration over several cycles of the underlying oscillation, yielding temporal smearing and uncertainty. These temporal correlations mean that decoding at one time point can be influenced by activity occurring slightly before or after that specific time—and this temporal dependence differs across EEG features. As a result, the temporal precision of our time-resolved RDMs differ across broadband scalp voltage and for frequency-band features (not to mention across different frequency bands), limiting the validity of cross-feature comparisons.

This caveat means that fine-grained timing differences–especially when comparing scalp voltage to band power–should be interpreted cautiously. Given these constraints, we focused our interpretation on effects that exceeded the temporal precision of our measures: scalp-voltage discriminability typically unfolded over windows of roughly 100 ms or more, and alpha-based discriminability was sustained over several hundred milliseconds, both longer than the temporal smearing introduced by preprocessing and time-frequency decomposition. Future work could systematically manipulate and match the temporal smoothing applied to different EEG features.

We trained classifiers for each feature independently for simplicity, but cross-feature interactions may influence results (Patel et al., 2022; Studenova et al., 2023). Interaction between different speech features such as talker location and talker identity modulate neural speech responses: spatial location affects mean response levels, while talker identity influences response variability (Patel et al., 2022; Zhang et al., 2016). Future work should therefore explore additivity across different channels and incorporate multi-feature interactions.

Beyond these limitations, RSA can be utilized in a broader context; to compare neural representations across brain regions, species, and imaging modalities. In particular, cross-modal RDMs could compare spatial and temporal representations between different modalities. Extending RSA to fMRI and magnetoencephalography (MEG) data revealed the spatiotemporal dynamics of object recognition and linking neural representations across time and space (Cichy et al. 2014). Salmela et al. (2016) also investigated distinct attention networks during audiovisual tasks by combining RSA with EEG- and fMRI-derived representations. This line of research could evolve into EEG-fMRI fusion RDMs, leveraging EEG’s temporal resolution and fMRI’s spatial specificity to provide insight into the spatiotemporal dynamics of attentional control across cortex (Cichy et al., 2020).

## Conclusions

This study showed that RSA can effectively capture both evoked and induced EEG responses during an auditory attention task. Scalp voltage signals reflected fast, cue-locked changes tied to auditory ERPs, while alpha power developed more slowly and was sustained throughout the task, closely tracking attentional demands. These features contributed differently over time, supporting their complementary roles in decoding attentional states. Overall, our findings support the use of RSA for uncovering the dynamics of attention in EEG data and suggest promising directions for future work, including cross-modal analyses with fMRI or MEG to better understand the spatial and temporal aspects of attentional processing.

## Acknowledgements

We thank Alex Pei for assistance with data collection, Tim Verstynen for helpful advice about the analysis, Vibha Viswanathan for valuable comments on the manuscript, and Maggie Henderson for suggesting the unique contribution analysis. This work was supported by the Office of Naval Research (N00014-53618-1-2069) and U.S. Department of Defense Hearing Restoration and Rehabilitation Program grants (HT9425-23-1-0912).

