## Supplementary Figure 2 for "Representational similarity analysis of EEG reveals multiple spatiotemporal dynamics of selective attention"

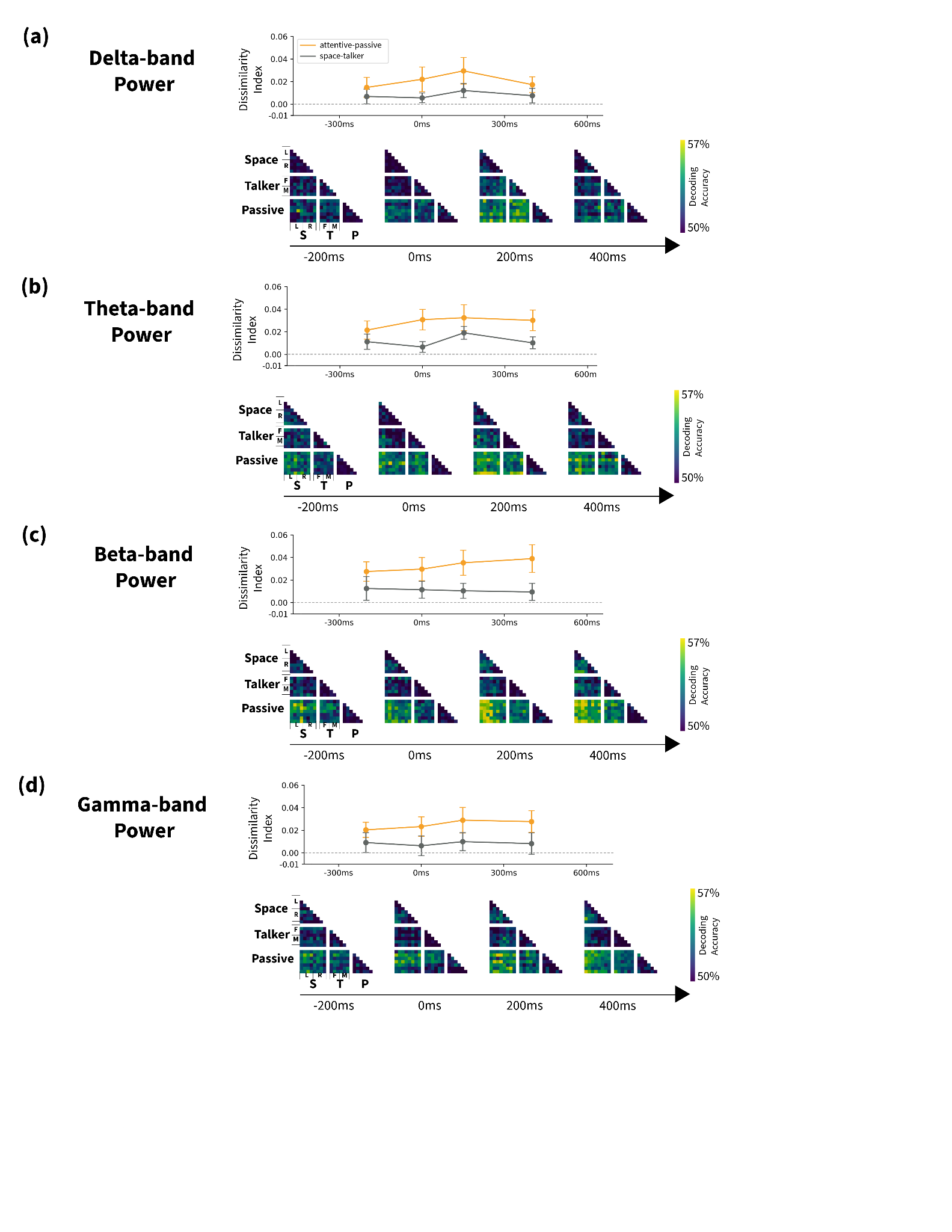

***Supplementary Figure 2. Representational dissimilarity during Stimulus Period in a) delta-, b) theta-, c) beta-, and d) gamma-band power.*** *Top: Dissimilarity indices in the representative time points for the contrast between attentive and passive listening (orange) and between spatial and talker attention (gray). Error bands are 95% within-subject, standard error-based confidence intervals of the mean. The horizontal dashed line indicates zero difference between the contrasted conditions. Bottom: Representational dissimilarity matrices (RDMs) at representative time points throughout the Cue Period.*
