## Supplementary Figure 5 for "Representational similarity analysis of EEG reveals multiple spatiotemporal dynamics of selective attention"

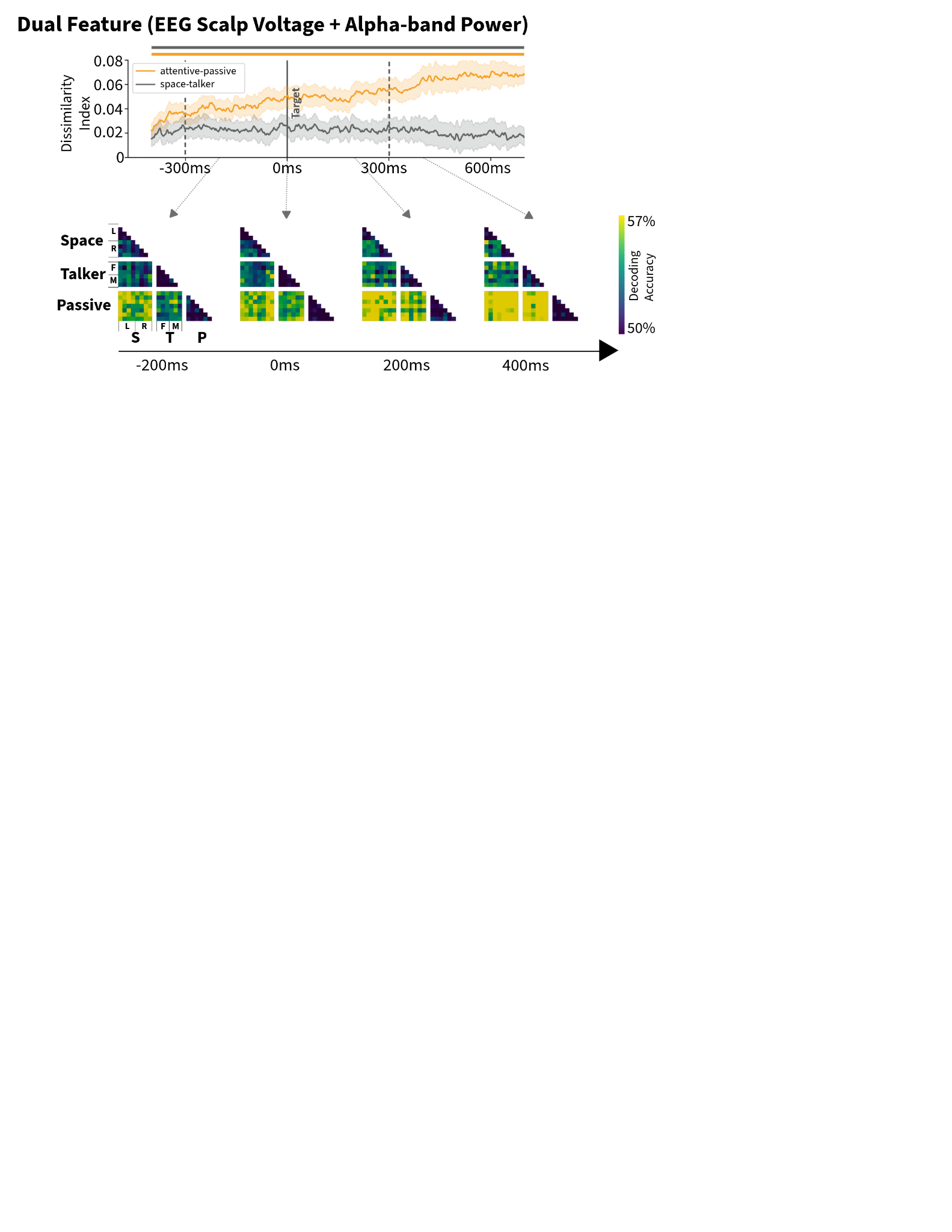

***Supplementary Figure 5. Representational dissimilarity during Stimulus Period using dual features, broadband scalp voltage and alpha-band power.*** *Top: Dissimilarity indices across the Cue Period for the contrast between attentive and passive listening (orange) and between spatial and talker attention (gray). Error bands are 95% within-subject, standard error-based confidence intervals of the mean. Onsets of the task cue (-1000 ms, dashed line) and target cue (0 ms, solid line) are indicated. Horizontal lines denote temporal clusters of statistical significance. Bottom: Representational dissimilarity matrices (RDMs) at representative time points throughout the Cue Period.*
