## Supplementary Figure 6 for "Representational similarity analysis of EEG reveals multiple spatiotemporal dynamics of selective attention"

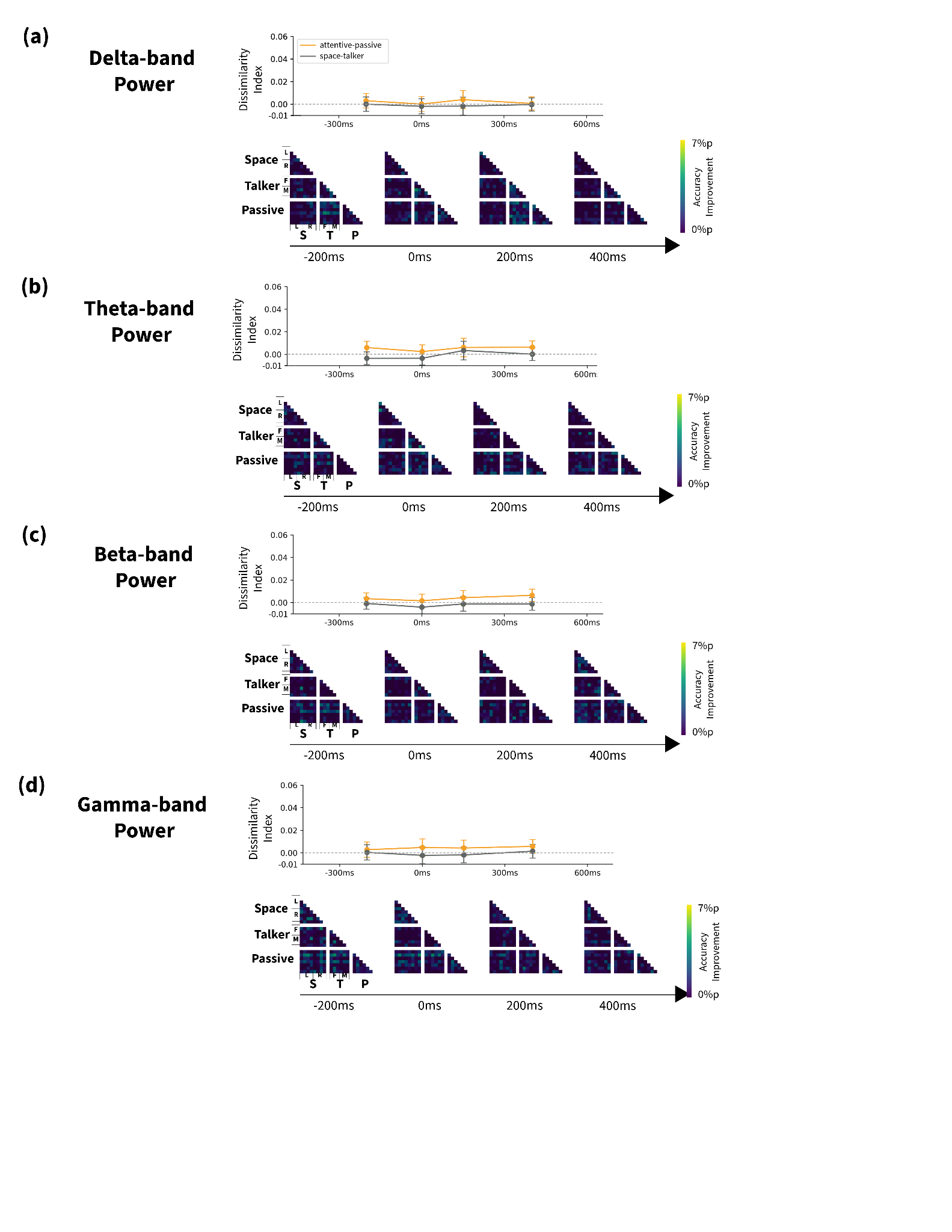

***Supplementary Figure 6. Unique contribution of a) delta-, b) theta-, c) beta-, and d) gamma-band power during Stimulus Period.*** *Top: Dissimilarity indices in the representative time points for the contrast between attentive and passive listening (orange) and between spatial and talker attention (gray). Error bands are 95% within-subject, standard error-based confidence intervals of the mean. The horizontal dashed line indicates zero difference between the contrasted conditions. Bottom: Representational dissimilarity matrices (RDMs) at representative time points throughout the Stimulus Period. RDM values show unique feature contributions, measured as the accuracy improvement (%p) of dual-feature decoding over each single-feature decoding.*
